# Exposure to a mixture of long-chain per- and polyfluoroalkyl substances (PFAS) disrupts ovarian follicle development, ovulation, and luteinization in female mice

**DOI:** 10.64898/2026.09.12.750977

**Authors:** Antonella R. R. Caceres, Jiyang Zhang, Tristan Costanza, Grace Guo, Qiang Zhang, Shuo Xiao

**Affiliations:** Department of Pharmacology and Toxicology, Ernest Mario School of Pharmacy, Rutgers University, Piscataway, NJ 08854, USA; Environmental and Occupational Health Sciences Institute (EOHSI), Rutgers University, Piscataway, NJ 08854, USA; Center for Environmental Exposures and Disease, Rutgers University, Piscataway, NJ 08854, USA; Gangarosa Department of Environmental Health, Rollins School of Public Health, Emory University, Atlanta GA 30322, USA

## Abstract

Per- and polyfluoroalkyl substances (PFAS) are persistent environmental contaminants associated with adverse female reproductive outcomes. However, most prior studies examined individual PFAS, despite that humans are exposed to complex PFAS mixtures. Here, we investigated the ovarian effects of a mixture of five commonly detected long-chain PFAS, including PFOS, PFOA, PFNA, PFDA, and PFHxS, using complementary *in vivo* mouse and *in vitro* 3D ovarian follicle culture models. Young adult female mice were exposed to a range of concentrations of the PFAS mixture (PFASmix) through daily drinking water for 8 weeks. PFASmix at high yet environmentally relevant concentrations (PFASmix-high) disrupted mouse estrous cyclicity, decreased circulating estradiol, increased testosterone, and promoted follicular atresia. Antral follicles from exposed mice showed decreased *Lhcgr* and increased *Amh* expression. Although the number of ovulated oocytes was unchanged following superovulation, PFASmix-high reduced oocyte size and polar body extrusion and altered oocyte cytoskeletal characteristics. In a 3D mouse ovarian follicle culture model, direct PFASmix exposure impaired follicle growth, ovulation, and oocyte maturation in a concentration-dependent manner. PFASmix-high suppressed the periovulatory expression of *Areg, Ereg, Tnfaip6,* and *Adamts1* and impaired corpus luteal (CL) development and function, as evidenced by reduced CL spheroid growth and cell survival, decreased expression of *Star, Cyp11a1, Hsd3b1,* and *Lhcgr*, and reduced progesterone secretion. PFASmix-high exposure additionally altered the expression of multiple PPARγ-associated genes in cultured ovarian follicles. Benchmark dose modeling identified estrous cyclicity and follicle rupture as sensitive *in vivo* and *in vitro* endpoints, respectively. Together, these findings demonstrate that exposure to the mixture of long-chain PFAS disrupts multiple stages of ovarian function, from follicular development, oocyte maturation to ovulation and luteal function, providing mechanistic and quantitative evidence for the reproductive toxicity of PFAS mixtures.

## Introduction

Per- and polyfluoroalkylated substances (PFAS) are a large group of synthetic chemicals that contain at least one fluorinated aliphatic carbon chain. Due to the high stability of the carbon-fluorine bond, since their first synthesis in the 1940s, PFAS have been widely used in consumer and industrial products, including non-stick coatings, food packaging, stain-resistant textiles, and aqueous firefighting film-forming foams (1). PFAS can be classified by their carbon chain length (long or short) and functional groups (carboxylic acids or sulfonates). Long-chain PFAS are composed of more than 6 or 8 carbon atoms in their backbone for sulfonic acids and carboxylic acids, respectively, including legacy PFAS such as perfluorooctanoic acid (PFOA), perfluooroctane sulfonic acid (PFOS), perfluorononanoic acid (PFNA), perfluorodecanoic acid (PFDA), and perfluorohexane sulfonic acid (PFHxS) (2).

PFAS have become an emerging environmental health concern due to their resistance to environmental and biological degradation, earning them the name of “forever chemicals” (3). The biological half-life of long-chain PFAS have been estimated at 8-15 years in humans (4, 5). Given their ubiquitous use and persistence, PFAS bioaccumulate in living organisms and biomagnify through the food chain (6, 7). Although the phase-out of several legacy PFAS, such as PFOA and PFOS, started over two decades ago (8), PFAS, particularly long-chain PFAS, are detectable in nearly more than 90% of populations, with blood concentrations ranging from 0.7 ng/mL in the general population to 1,725 ng/mL among fluorochemical workers (9–15). In addition to systemic circulation, PFAS have also been reported to distribute to various organ systems, including the kidneys, brain, liver (16), bones (17), reproductive tissues and fluids (18–20).

Exposure to PFAS is associated with various harmful health effects, including hepatotoxicity (21, 22), carcinogenesis (23, 24), altered immune response (25), thyroid dysfunction (26), and disrupted lipid metabolism (27). The underlying mechanisms might involve interactions between PFAS and nuclear receptors, including the peroxisome proliferator-activated receptor (PPAR), estrogen receptor, and thyroid hormone receptor (28–30). Accumulating evidence indicates that PFAS act as endocrine-disrupting chemicals (EDCs) to impair female reproduction (4, 31, 32). In humans, exposure to PFAS has been related to primary ovarian insufficiency (POI) (33, 34), irregular menstrual cycles (35, 36), polyendocrine metabolic ovarian syndrome (PMOS, previously termed as polycystic ovary syndrome or PCOS) (37, 38), and infertility (39, 40). Experimental research reported adverse effects of PFAS on the hypothalamic-pituitary-ovarian (HPO) axis (41), ovarian cyclicity, steroidogenesis, and oocyte maturation (42–44).

Despite the large amount of evidence linking PFAS exposure to female reproductive disorders, most prior studies, however, only examined individual PFAS, which did not reflect the real-world scenario of human exposure to PFAS mixtures. Exposure to PFAS mixture may result in significantly different and potentially more severe health effects than individual PFAS, as co-occurring PFAS compounds may act through divergent, overlapping, or shared molecular pathways (29). Our previous study demonstrated that individual long-chain PFAS, including PFOA, PFOS, and PFNA, disrupt gonadotropin-dependent follicle development, steroidogenesis, and ovulation, whereas the tested short-chain PFAS produced substantially weaker ovarian effects (42). Mechanistically, we identified PPARγ activation in follicular granulosa cells as a molecular initiating event (MIE) for PFNA-induced ovarian toxicity. Because multiple long-chain PFAS can interact with PPARs and are commonly present together in human and environmental samples, these findings raise an important question yet to be answered: does exposure to PFAS mixture disrupt ovarian function, and does PPAR-initiated signaling mediate a convergent molecular response?

In the present study, we hypothesize that exposure to an environmentally relevant PFAS mixture impairs female ovarian functions and related reproductive outcomes, potentially via altered PPAR signaling pathways. To test this hypothesis, we conducted an *in vivo* study by exposing young adult female mice to real-world levels of PFAS mixture in drinking water and examining the effects on ovarian functions. Further, we employed a 3D *in vitro* mouse ovarian follicle culture system to assess the direct ovarian effect of PFAS mixtures and the underlying toxic mechanisms.

## Materials and Methods

### Animals

CD-1 mice were purchased from Envigo (Indianapolis, IN) and maintained at the Animal Care Facility in the Research Tower at Rutgers University. The animals were housed in polypropylene cages in a temperature (22±1°C), humidity (30%–70%), and light/dark cycle (12/12) controlled environment, with food (PicoLab Mouse Diet 20 EXT 5R58, Cat. No.3003269-712; LabDiet) and RO (reverse osmosis) water available *ad libitum*. Sixteen-day-old female mice were used for preantral follicle isolation and *in vitro* culture. Eight-week-old female mice were used for *in vivo* oral PFAS exposure. All experimental procedures were reviewed and approved by the Institutional Animal Care and Use Committee (IACUC) at Rutgers University.

### PFAS mixture composition and selection of exposure concentrations

Five long-chain PFAS, including PFHxS, PFNA, PFOS, PFOA, and PFDA, were selected to construct the PFAS mixtures, representing legacy long-chain PFAS frequently detected in human biomonitoring studies, including the 2017–2020 National Health and Nutrition Examination Survey (NHANES) (45). Information on the chemical identity, classification, CAS number, and purity of each PFAS is provided in Supplementary Table 1.

For the *in vivo* exposure study, the relative composition of the five-PFAS mixture was based on groundwater contamination data from Marine Corps Base Camp Lejeune, North Carolina, obtained from the Naval Facilities Engineering Systems Command (NAVFAC) groundwater investigation and the PFAS Exchange contamination database (46, 47) and can be found in Supplementary Table 2. Four mixture concentrations, hereafter referred to as PFASmix-low, -medium, -medium-high, and -high, were evaluated while maintaining the same concentration ratio of the five PFAS compounds. PFASmix-low was based on the upper limit of total concentration of the five PFAS compounds identified in a U.S. public water system in the EPA Fifth Unregulated Contaminant Monitoring Rule (UCMR 5) dataset (48). PFASmix-medium was based on the selected Camp Lejeune groundwater contamination profile, and PFASmix-medium-high and PFASmix-high were 10- and 100-fold higher than PFASmix-medium, respectively. This concentration range was designed to span concentrations documented in contaminated drinking water and groundwater, including highly contaminated sites, while enabling characterization of the dose-response relationship. The total PFAS mass concentrations for PFASmix-low, -medium, -medium-high, and - high were 4.34 × 10⁻□, 4.35 × 10⁻□, 4.35 × 10⁻□, and 4.35 × 10⁻³ mg/mL, respectively (Supplementary Table 3).

For the *in vitro* study, the relative composition of the five-PFAS mixture in the follicle culture medium was based on human serum concentrations (Supplementary Table 4), which are a more appropriate internal exposure reference than drinking-water concentrations. We tested five PFASmix concentrations, with the lowest concentration, also designated as PFASmix-low, anchored on the geometric mean serum concentrations of PFHxS, PFNA, PFOS, PFOA, and PFDA reported in NHANES 2017–2020 (49). The four higher mixture concentrations are sequentially 10, 100, 1,000, and 10,000-fold higher than PFASmix-low, while maintaining the same concentration ratio between the five PFAS (Supplementary Table 5). The resulting total PFAS mass concentrations were 9.75 × 10⁻□, 9.75 × 10⁻□, 9.75 × 10⁻□, 9.75 × 10⁻³, and 9.75 × 10⁻² mg/mL, corresponding to molar concentrations of 0.021, 0.21, 2.1, 21, and 210 µM, respectively. The highest concentration of 210 µM is justified because previous human biomonitoring studies have reported PFAS serum concentrations spanning several orders of magnitude, with PFOA and PFOS reaching approximately 222 µM in occupationally exposed populations (5, 50–52).

Individual PFAS stock solutions were prepared in dimethyl sulfoxide (DMSO; Sigma-Aldrich). PFASmix working solutions were freshly prepared by dilution of the stock solutions in RO water for *in vivo* exposure or in culture medium for *in vitro* exposure. The final DMSO concentrations were 0.00001% in drinking water and 0.01% in follicle culture medium. Vehicle controls contained equivalent concentrations of DMSO.

### *In vivo* PFAS mixture exposure and sample collection

Animals were fed with drinking water containing the PFASmix for 8 weeks (Figure 1A). A total of 45 mice were randomly assigned to the control (vehicle DMSO) or PFASmix treatment groups. Mouse food and water consumption, body weight, and other general health endpoints, such as survival and behavior, were examined daily and summarized weekly. Water was replenished once the volume fell below ∼ 50 mL. Vaginal smears were performed daily in the last 3 weeks of the exposure to assess estrous cycles. At the end of the 8^th^ week, animals were euthanized in the morning of the proestrus with CO_2_ asphyxiation, followed by decapitation. For animals with a disrupted estrous cycle in which proestrus could not be identified in the 8^th^ week, they were sacrificed on the last day of the 8^th^ week. Upon animal sacrifice, trunk blood was immediately collected and centrifuged at 16,000 × g for 15 min to separate serum for hormone measurements. Ovaries were harvested, weighed, and either immediately fixed in 4% paraformaldehyde (PFA) for histology or snap-frozen for total RNA extraction and RT-qPCR. One of the two ovaries from each mouse was kept in dissection media (L15 media, 0.5% pe-strep, 1% FBS) and further dissected to obtain large antral follicles (diameter >300µm), which were then snap-frozen in Picopure® extraction buffer for RT-qPCR.

**Figure 1.**
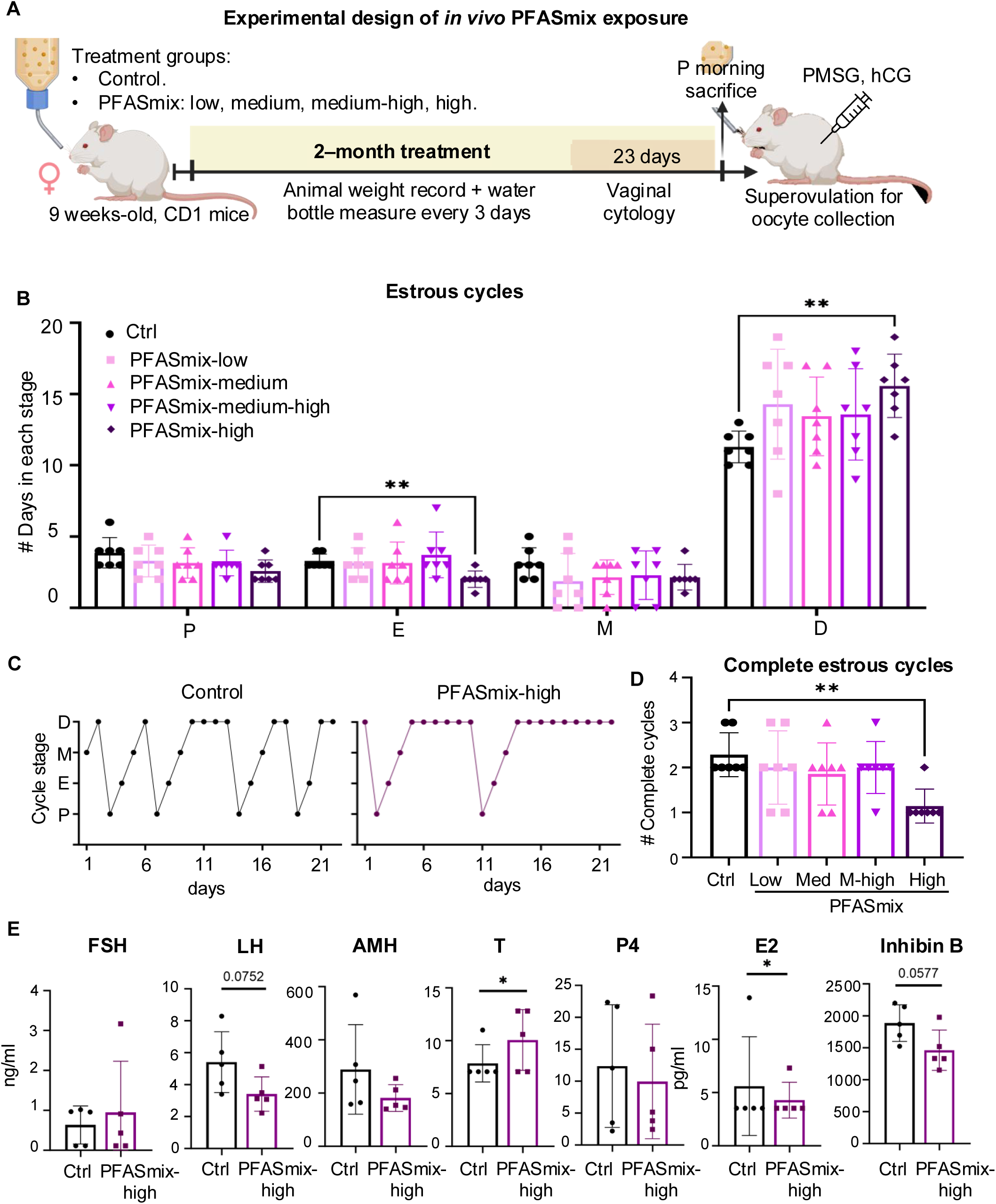
*In vivo* experimental design, estrous cyclicity, and serum hormone concentrations after two months of PFAS mixture exposure in drinking water. (A) Schematic for the *in vivo* experimental design. (B) Number of days at each stage of the estrous cycle (P: proestrus, E: estrous, M: metestrus, and D: diestrus) through a span of 23 days in control (Ctrl), low, medium (Med), medium-high (Med-high), and high PFAS mixture (PFASmix) concentration. (C) Estrous cycle overview of a control and a PFASmix-high-exposed animal. (D) Number of complete estrous cycles (from estrus to estrus) registered for each animal during the 23-day period. (E) Serum hormone concentration is measured in ng/ml for FSH, LH, AMH, testosterone (T), and progesterone (P4), and in pg/ml for estradiol (E2) and inhibin-b. PMSG: Pregnant mare’s serum gonadotropin, hCG: Human chorionic gonadotropin. Bars represent mean ± SD, asterisks represent significant differences between the connected groups. *\*p<0.05; **p<0.01*.

### Superovulation

To evaluate the effects of the PFASmix on ovulation *in vivo*, 10 animals were exposed to either vehicle or PFASmix-high for 8 weeks. Mice received an intraperitoneal (i.p.) injection of 5 IU of pregnant mare serum gonadotropin (PMSG, ProSpec), followed by another i.p. injection of 5 IU of human chorionic gonadotropin (hCG, Sigma-Aldrich) 46 hours later to induce ovulation. Animals were sacrificed 16 hours after the hCG injection, and the ovaries and oviducts were collected in dissection media. The cumulus-oocyte complexes (COCs) in the oviducts were examined and counted under a stereoscope after puncturing the oviduct ampulla. The oocytes were enzymatically denuded using a 0.3% hyaluronidase solution (Sigma-Aldrich), imaged for classification, and fixed for immunofluorescence analysis.

### Follicle culture, development, ovulation, and luteinization in the *in vitro* 3D culture

Ovaries of 16-day-old CD1 mice were collected and processed to obtain multilayered secondary follicles as described previously (53). Briefly, mice were euthanized with CO_2_ asphyxiation followed by cervical dislocation. The ovaries were harvested in dissection media and, after mechanical and enzymatic isolation, further dissociated into single follicles. Secondary follicles with a diameter of 150-180 µm were selected and encapsulated in 0.5% alginate hydrogel. Individual follicles were cultured in 96-well plates with 100 µl of growth media (50% αMEM Glutamax, 50% F-12 Glutamax, supplemented with 3 mg/ml BSA, 5 mIU/ml hFSH [Gonal-F®], 1 mg/ml bovine fetuin, and 5 µg/ml ITS) at 37°C and 5% CO_2_. Follicles were allowed to grow in basal growth medium for 48 hours; thereafter, they were divided into vehicle- or PFASmix-treated groups.

Follicle growth was measured every 2 days before media change. After 6 days of growth, the follicles were released from the alginate encapsulation by incubation in L15 media containing 1% FBS and 10 IU/mL alginate lyase (Sigma Aldrich) at 37 °C for 10 minutes. At this stage, some follicles were collected to assess expression of marker genes for follicular development, and the conditioned media were collected for subsequent hormone analysis by ELISA. The remaining follicles were transferred to a new 96-well plate with 100 µl of ovulation media (same composition as growth media, plus 1.5 IU/mL hCG (Sigma-Aldrich) and 10 mIU/mL hFSH), and each experimental group was treated with the corresponding vehicle or PFAS mixture concentration. After 4 hours of ovulation induction, a set of follicles was collected to evaluate ovulation-related gene expression (53). At 16 hours post-hCG induction, follicle rupture or “ovulation” was assessed under an Olympus inverted microscope with a 10x objective (Olympus Optical Co Ltd, Tokyo, Japan) using Tcapture imaging software (Tucsen, V5.1.1). A follicle was classified as ruptured when one side of the follicular wall was breached, and unruptured when the wall was intact. The oocytes were collected and morphologically classified as follows: oocytes with extruded polar bodies were classified as metaphase II (MII), and all others (germinal vesicle or germinal vesicle breakdown) were considered as non-MII. After classification, oocytes were fixed and maintained in blocking buffer until further processing. The follicles devoid of oocytes after induced ovulation were cultured for 48 hours in fresh ovulation induction media without hFSH to induce luteinization and progesterone (P4) secretion. Luteal spheroids were then imaged, collected, and fixed in 4% PFA for histological assessment or frozen for RT-qPCR. The conditioned media were saved for P4 ELISA determination. Follicles were imaged at each media change every other day, using the 10x objective of the Olympus inverted microscope. Follicle diameter was determined by averaging the lengths of two perpendicular measurements using ImageJ software (v1.53; National Institutes of Health, NIH, Bethesda, MD).

### Hormone measurements

Serum aliquots were sent to the University of Virginia (UVA) Center for Research in Reproduction Ligand Assay and Analysis Core. FSH and LH were measured with in-house ultra-sensitive ELISAs based on the methods published by Ongaro et al. (54) and Steyn et al. (55). The remaining hormones were measured using commercially available ELISA kits, following the manufacturers’ instructions: estradiol (E2, ALPCO, 11-ESTHU-E01); AMH and inhibin B (ANSH LAB, AL-113, AL-163); P4 and testosterone (T) (IBL, IB79105, IB79106). Each assay characteristic is summarized in Supplementary Table 6.

T, E2, and P4 concentrations in conditioned media were measured using ELISA kits (Cayman Chemical, Ann Arbor, MI; catalog numbers 582701, 501890, and 582601, respectively) according to the manufacturer’s instructions. The absorbance was measured using a BioTek SpectraMax M3 microplate reader (BioTek Instruments, Inc., Winooski, VT) at the recommended wavelengths. Reportable ranges for each assay were: 0.61-10,000 pg/ml for E2, 3.9-500 pg/ml for T, and 7.8-1,000 pg/ml for P4. The sensitivities were 20, 6, and 10 pg/mL for E2, T, and P4, respectively. Inter-assay variability (CV) for E2, T and P4, was 8%-12.3%, 7.7%-14.2% and 7.7%-16.4%; intra-assay CV was 6.5%-10.8%, 4.4%-19.1% and 7.3%-14.5%.

### RNA extraction and RT-qPCR

For *in vivo* animal exposure, large antral follicles (>300 µm) were manually dissected and saved in extraction buffer at −80 °C until further processing. For *in vitro* experiments, follicles were extracted from alginate beads using alginate lyase and were stored in extraction buffer at −80 °C until further processing, along with CLs. Total RNA was extracted using the Arcturus™ PicoPure™ RNA Isolation Kit (Thermo Fisher Scientific). RNA quantification and the A260/A280 ratio were assessed with a NanoDrop. RNA was then reverse-transcribed into cDNA using the Superscript Strand Synthesis System with random hexamer primers (Invitrogen) and stored at −80 °C. RT-qPCR was performed in 384-well plates using the Power SYBR Green PCR Master Mix (Thermo Fisher Scientific) in a ViiA 7 Real-Time 436 PCR System (Thermo Fisher Scientific). The thermocycler was programmed for 10 min at 95 °C, followed by 40 cycles of 15 seconds at 95 °C and 40 seconds at 60 °C, and concluded with a melting curve to determine primer specificity. Relative expression levels of each gene were normalized by the expression of β-actin (Actb). For *in vitro* and *in vivo* exposure studies, 8 follicles were included per treatment group. The primer sequences used are listed in Supplementary Table 7.

### Histology evaluation

Tissue fixation was achieved by incubation in 4% PFA at 4□°C overnight. Once fixed, tissues were washed in PBS and dehydrated in ethanol washes before xylene clarification and paraffin embedding. For processing luteal spheroids generated *in vitro*, the structures were first embedded in a 0.5% alginate hydrogel prior to fixation. The tissues were then sectioned to 5 µm thickness and stained with hematoxylin and eosin for histopathological assessment.

Ovaries were serially sectioned at 5 μm, and follicles at different developmental stages were counted every 10 sections. Follicles were classified as primordial follicles (containing an oocyte surrounded by flattened pre-granulosa cells), primary follicles (containing an oocyte surrounded by a single layer of cuboidal granulosa cells), secondary follicles (containing an oocyte surrounded by > 1 granulosa cell layer), and antral follicles (containing an oocyte surrounded by multiple layers of granulosa cells with an antral cavity and a theca cell layer). Atretic follicles (degenerated follicles with high granulosa cell apoptosis, fragmented oocyte, and thickened theca layer) and CL (structures formed by luteal cells with characteristic cytoplasmic eosinophilia) were also counted.

### Oocyte immunofluorescence

Oocytes collected from the *in vivo* superovulation experiment and from *in vitro* ovulation were fixed in 2% PFA for 20 minutes at room temperature and saved in blocking buffer (PBS, 0.3% BSA, 0.01% Tween-20, and 0.02% NaN3) at 4 °C until processed. The oocytes were then permeabilized in PBS containing 0.3% BSA, 0.01% Triton X-100, and 0.02% NaN3 for 15 minutes at room temperature, followed by 3 washes in blocking buffer. Immunostaining was performed by incubating the oocytes in a solution containing the primary antibodies Rhodamine phalloidin (Invitrogen, R415, for F-actin staining) and α-tubulin (Cell Signaling, 11H10, rabbit monoclonal conjugated with Alexa Fluor 488) at a 1:100 dilution in blocking buffer for 1 hour in a dark, humidified chamber at 4 °C. The cells were then washed with blocking buffer and mounted with 10 μL of VectaShield (Vector Laboratories Inc., Newark, CA).

### *In situ* RNA hybridization

Ovaries from *in vivo* superovulation (containing new and old CL) and luteal spheroids collected on day 9 of *in vitro* incubation were fixed, paraffin-embedded, and sectioned to 5 µm thickness. To detect RNA transcripts *in situ*, the RNAscope Multiplex Fluorescent Detection Kit V2 and the HybEZ Hybridization System (Advanced Cell Diagnostics, Newark, CA) were used. In accordance with the manufacturer’s instructions, tissue sections were prepared via heat-induced epitope retrieval, hydrogen peroxide treatment, and protease-based permeabilization. Following the hybridization of gene-specific probes, the signal was amplified using horseradish peroxidase (HRP) and visualized with fluorescent dyes. Finally, sections were mounted in DAPI-enriched Vectashield antifade medium and imaged using a Leica confocal microscope (Wetzlar, Germany).

### TUNEL assay

Apoptotic detection was performed using the DeadEnd™ Fluorometric TUNEL System (Promega, Madison, WI) according to the manufacturer’s protocol. Briefly, the slides were heated, the paraffin was removed with xylene washes, and they were rehydrated for apoptosis detection. The samples were fixed with 4% PFA in PBS and permeabilized with 20µg/ml proteinase K solution. Then the slides were equilibrated with the equilibration buffer provided and labeled with the TdT reaction mix for 60 minutes at 37°C in a humidified chamber. The label reaction was stopped by 2x SCC incubation, and the slides were washed 3 times with a PBS solution containing 0.1% Triton X-100 and 5 mg/ml BSA to reduce background noise. Finally, sections were mounted in DAPI-enriched Vectashield antifade medium and imaged in a confocal fluorescence microscope.

### Benchmark dose/concentration modeling

Dose or concentration-response relationships were evaluated using the EPA Benchmark Dose Software Online (BMDS Online). Benchmark concentration (BMC) modeling was performed for selected *in vivo* estrous-cycle endpoints and *in vitro* follicle endpoints showing concentration-dependent responses. For continuous endpoints, a benchmark response (BMR) corresponding to a 5% relative deviation from the control mean (BMR_5_) was used. Follicle non-rupture was analyzed as a dichotomous endpoint using a BMR of 10% extra risk (BMR_10_). The best-fitting model was selected according to BMDS model-fit criteria. BMC estimates, and corresponding lower and upper confidence bounds (BMCL and BMCU) were obtained for each endpoint.

### Statistical analysis

Statistical analysis was performed using GraphPad Prism (version 10; GraphPad Software). Two-way ANOVA followed by Dunnett’s multiple comparisons test was used to compare numerical data across treatment groups, including the number of days spent in each estrous cycle stage, the number of ovarian structures per section, and RT-qPCR gene expression data (in vivo follicle genes, liver genes, in vitro follicle maturation and ovulation-related genes, corpus luteum genes, and PPARγ-related genes). The number of complete estrous cycles was analyzed by the Kruskal-Wallis test followed by Dunn’s post hoc test. One-way ANOVA followed by Dunnett’s post hoc test was used for in vitro follicle size. In vitro estradiol, testosterone, and progesterone concentrations were log-transformed to address skewed distribution and analyzed by one-way ANOVA followed by Dunnett’s post hoc test. Unpaired Student’s t-test was used to compare two-group numerical data, including serum AMH, progesterone, inhibin-B, and LH concentrations; oocyte diameter (*in vivo* and *in vitro*); in vivo spindle fluorescence intensity and actin cap width; liver weight; in vitro spindle volume and fluorescence intensity; corpus luteum cell number (*in vitro*); and uterine receptor gene expression. The Mann-Whitney rank-sum test was used for non-normally distributed two-group data, including serum testosterone, estradiol, and FSH concentrations; in vivo oocyte number; in vivo spindle volume; in vitro actin cap width; and corpus luteum diameter. Fisher’s exact test was used to analyze categorical data, including in vivo follicle TUNEL labeling, in vivo oocyte maturation classification, in vitro follicle rupture percentage, polar body extrusion, oocyte maturation rate, and in vitro corpus luteum TUNEL labeling. Data are reported as mean ± SD. Statistical significance was defined as p<0.05.

## Results

### *In vivo e*xposure to an environmentally high yet human-relevant level of PFASmix altered mouse estrous cycles

Young adult female mice were exposed to vehicle or increasing concentrations of PFASmix via drinking water for 8 weeks to examine their reproductive effects (Figure 1A). Water consumption and body weight gain were comparable across treatment groups, and no treatment-related changes in survival or overt clinical signs were observed during the exposure period. Daily vaginal cytology during the final 3 weeks of exposure showed that PFASmix-low, -medium, and -medium-high doses did not significantly alter the estrous cyclicity. In contrast, oral exposure to PFASmix-high significantly increased the number of days in diestrus (*p = 0.0027*), decreased the number of days in estrus (*p=0.0047*), and reduced the number of complete estrous cycles within the 3 weeks (*p=0.0226*; Figure 1B–D).

### Exposure to PFASmix-high altered mouse circulating levels of ovarian hormones in vivo

Based on the disruption of estrous cyclicity, we selected PFASmix-high for further evaluation of reproductive endpoints. Oral exposure to PFASmix-high via daily drinking water for 8 weeks did not significantly alter circulating FSH, AMH, or P4 concentrations but significantly decreased E2 (*p=0.0476*) and increased T (*p=0.0317*) (Figure 1E). LH (*p=0.0752*) and inhibin B (*p=0.0577*) showed decreasing trends that did not reach statistical significance.

### Exposure to PFASmix-high induced mouse follicle atresia *in vivo*

Histological staining and follicle counting results showed that after 2-month oral exposure, the ovaries from mice treated with PFASmix-high had comparable numbers of primordial, primary, secondary, and antral follicles to the control group (Figure 2A-B). However, PFASmix-high significantly increased the number of atretic follicles (*p=0.0469*). Notably, most atretic follicles were antral follicles, with TUNEL staining indicating significantly more apoptotic cells in the mural granulosa cell layers (*p<0.0001*) (Figure 2C-D).

**Figure 2.**
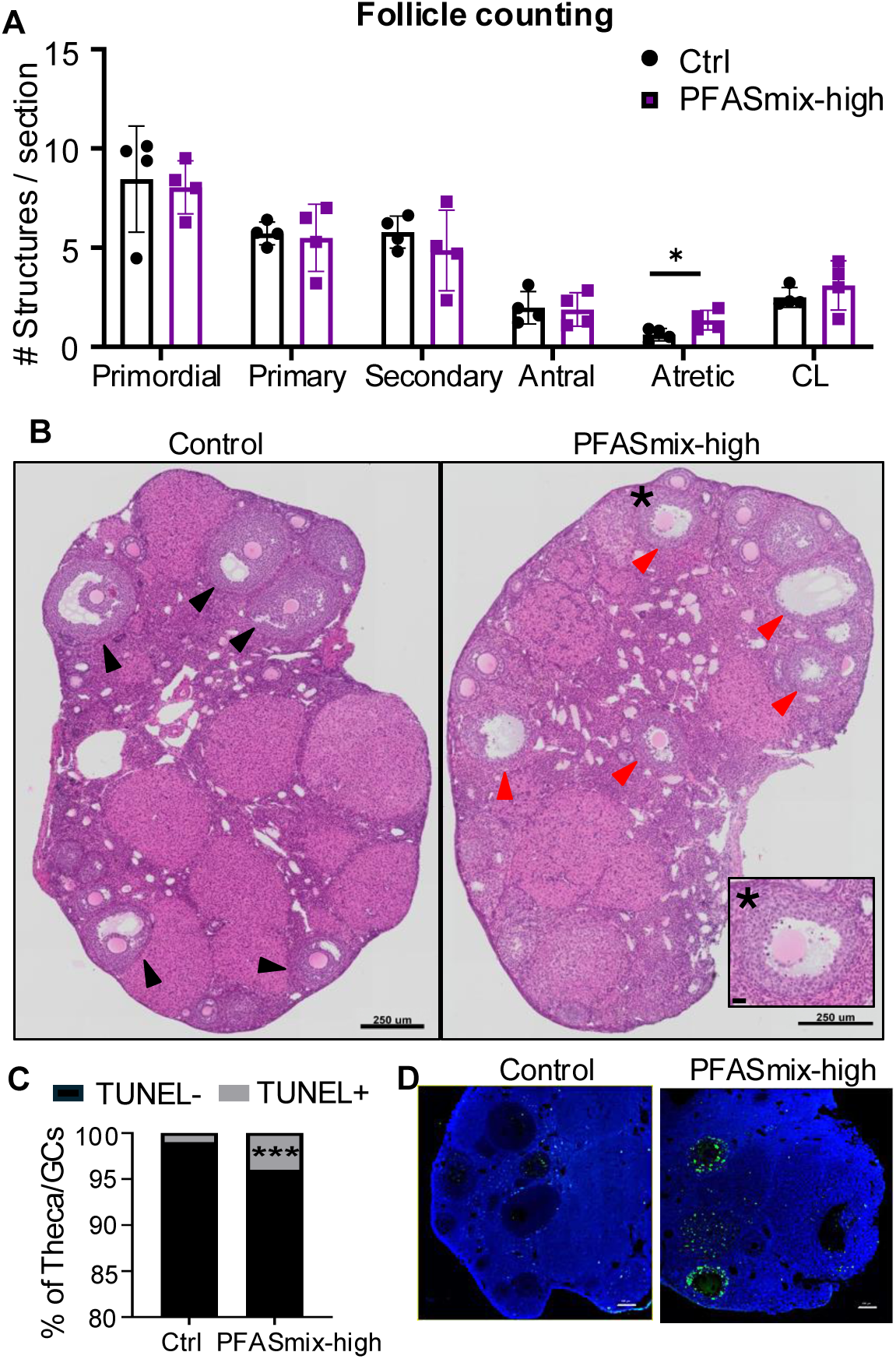
*In vivo* ovarian follicle counting and ovarian terminal deoxynucleotidyl transferase dUTP nick-end labeling (TUNEL) stain. (A) Number of ovarian structures per section in control and PFASmix-high exposure. (B) Representative photomicrograph of a control (Ctrl) and a PFASmix-high-treated animal’s ovary. Black arrowheads point to antral follicles. Red arrowheads point to atretic antral follicles, and an asterisk marks the zoomed-in atretic follicle in the lower-right corner. Scale bar 250 µm, inset scale bar: 20 µm. (C) Percentage of theca and granulosa cells labeled with TUNEL in control and PFASmix-high exposed animals. (D) Representative photomicrograph from an ovary from a control animal and a PFASmix-high exposed animal, after TUNEL stain. CL: corpora lutea. Scale bar 250 µm*. *p<0.05, ***p<0.0001*.

### Exposure to PFASmix-high reduced the expression of follicle development marker genes

The reduced serum concentrations of E2 and Inhibin B suggest that PFAS may interfere with gonadotropin-dependent follicle development. We next isolated large antral follicles (>300 µm) from control and PFASmix-high-treated mice on proestrus and analyzed gene expression by RT-qPCR. PFASmix-high significantly decreased *Lhcgr* expression (*p=0.000228*) and increased *Amh* expression (*p=0.000934*), whereas other genes related to follicular development were not significantly altered (Figure 3A-B). RNAscope analysis of ovarian sections further verified the reduced *Lhcgr* in granulosa cells of antral follicles from PFASmix-high-treated mice, as well as in theca cells of growing follicles (Figure 3C).

**Figure 3.**
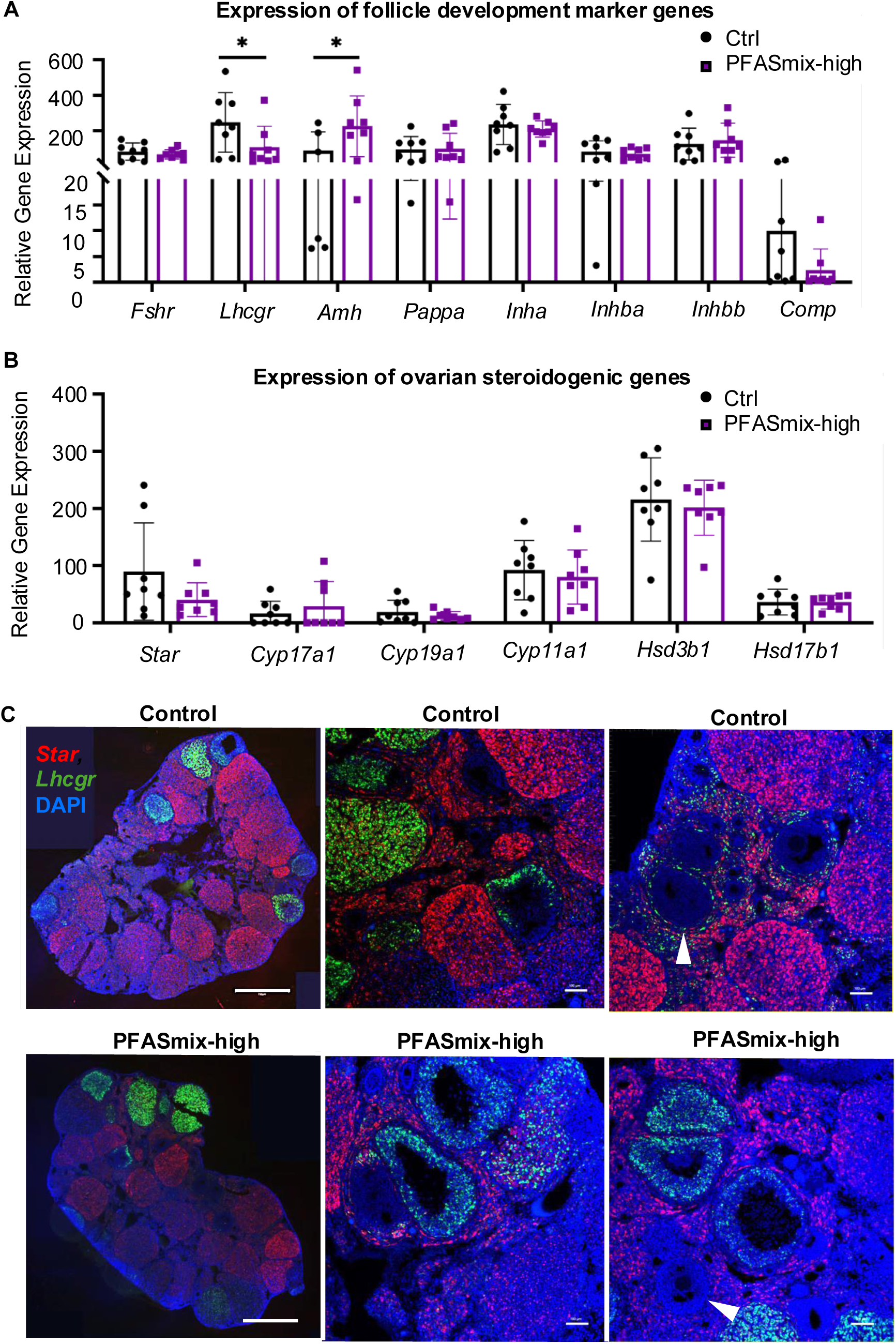
*In vivo* antral follicle expression of development-related genes and whole ovary RNAscope. (A) Relative gene expression of follicle development genes and (B) steroidogenic-related genes was examined by RT-qPCR in single antral follicles obtained from control (Ctrl) and PFASmix-high 2-month-exposed animals. The mRNA expression levels were normalized to β-actin expression. *p<0.05. (C) RNAscope representative images of control (top row) and PFASmix-high-exposed (bottom row) animals after superovulation, showing DAPI (blue), *Star* (red), and *Lhcgr* (green). White arrowheads point to theca cells in control and PFASmix-high antral follicles. Scale bars: 100 µm for zoomed-in pictures, and 750 µm for the whole ovary image. *\*p<0.05*.

### PFASmix-high impaired oocyte maturation and cytoskeletal characteristics *in vivo*

Following 8 weeks of exposure, control and PFASmix-high-treated mice underwent superovulation induction, and ovulated oocytes were recovered from the oviducts for morphological and cytoskeletal analyses. PFASmix-high did not alter the number of ovulated oocytes (Figure 4A) but significantly reduced the percentage of oocytes undergoing polar body extrusion (*p < 0.0071*; Figure 4B) and the oocyte size (*p=0.0181*; Figure 4C). Immunofluorescence analysis showed no significant change in meiotic spindle volume (*p=0.3031*) but revealed reduced spindle fluorescence intensity (*p=0.0019*) and reduced actin-cap width in oocytes from PFASmix-high-treated mice (Figure 4D–4G).

**Figure 4.**
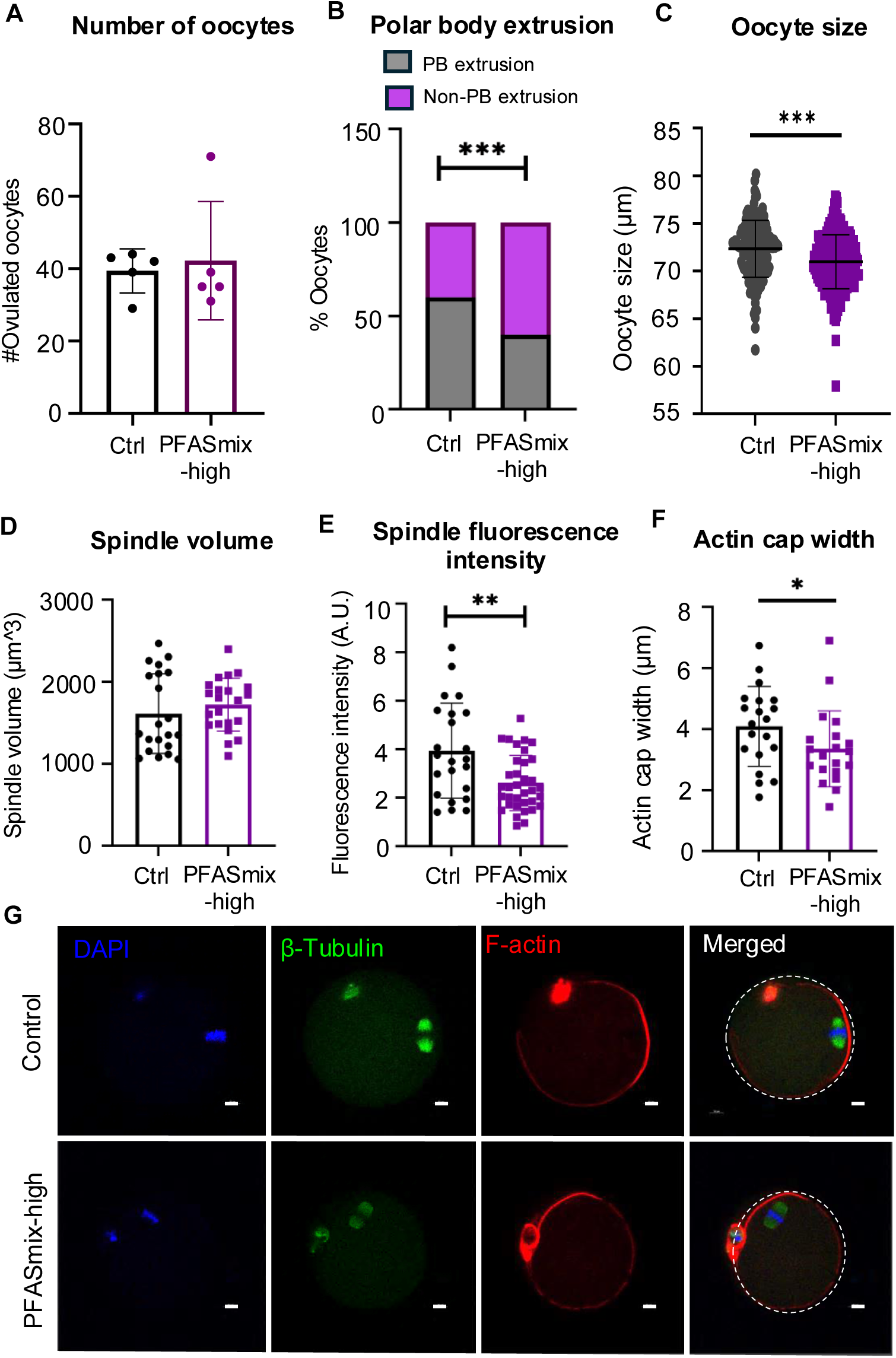
*In vivo* superovulation results and oocyte morphology. (A) The number of ovulated oocytes that were collected from the ampulla after superovulation treatment, and their (B)polar body extrusion rate in control (Ctrl) and PFASmix-high exposed animals. (C) Oocyte size (diameter, µm) of ovulated oocytes. (D) Meiotic spindle volume (µm³) and spindle fluorescence intensity, measured from β-tubulin immunofluorescence in control and PFASmix-high oocytes. (F) Actin cap width (µm) in control and PFASmix-high oocytes. (G) Representative confocal images of ovulated oocytes from control (top row) and PFASmix-high (bottom row) animals stained for DNA with DAPI, for the meiotic spindle with anti-β-tubulin, and for F-actin, together with the merged channel; dashed outlines in the merged panels indicate the oocyte membrane. Scale bars: 10 µm. *\*p<0.05, **p<0.01 and ***p<0.001*.

### PFASmix inhibits follicle development, ovulation, and oocyte maturation *in vitro*

We next used our established 3D *in vitro* follicle growth system, IVFG, to better understand the direct effect of PFASmix on gonadotropin-dependent follicle development and ovulation. Immature mouse follicles were treated with various concentrations of PFASmix during gonadotropin-dependent phase of follicle development and hCG-stimulated ovulation *in vitro* (Figure 5A). After ovulation induction, we continuously exposed formed CL spheroids to PFASmix for 2 days to assess CL morphology and function. The results showed that *in vitro* exposure to PFASmix-high significantly inhibited follicle growth starting from day 4 of IVFG (*p=0.0003*), resulting in smaller follicles by day 6 (*p<0.0001*) (Figure 5B-5C). Following ovulation induction, PFASmix concentration-dependently inhibited follicle rupture and oocyte polar body extrusion (Figure 6A-5C).

**Figure 5.**
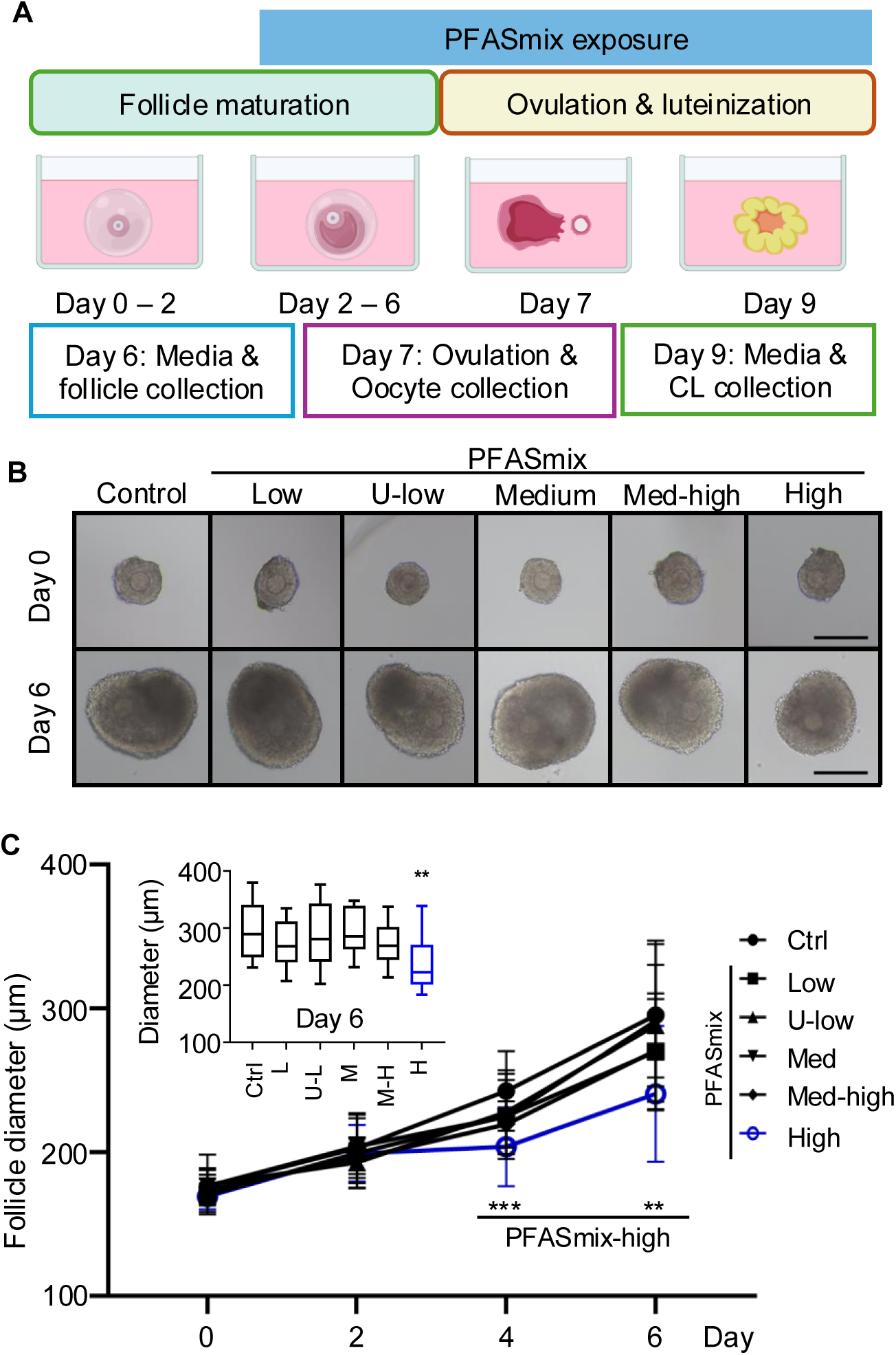
*In vitro* experimental design using *in vitro* follicle growth (IVFG) and effects of PFAS mixture exposure on follicle growth. (A) Schematic of the IVFG experimental design. Follicle PFAS mixture exposure started from day 2 of culture and was replenished every media change onwards. Conditioned media and follicles were collected on day 6, oocytes were collected on day 7 following ovulation induction, and media and CL (corpus luteum) samples were collected on day 9. (B) Representative brightfield images of encapsulated follicles at day 0 (top row) and day 6 (bottom row) of culture for the control (Ctrl), low, upper-lower (U-low), medium (Med), medium-high (Med-high), and high PFAS mixture concentration groups. Scale bars: 200 µm. *\*\*p<0.01 and ***p<0.001.* (C) Follicle diameter from IVFG cultured day 0 to day 6, with inserted box plot showing the diameters on day 6 across all groups.

**Figure 6.**
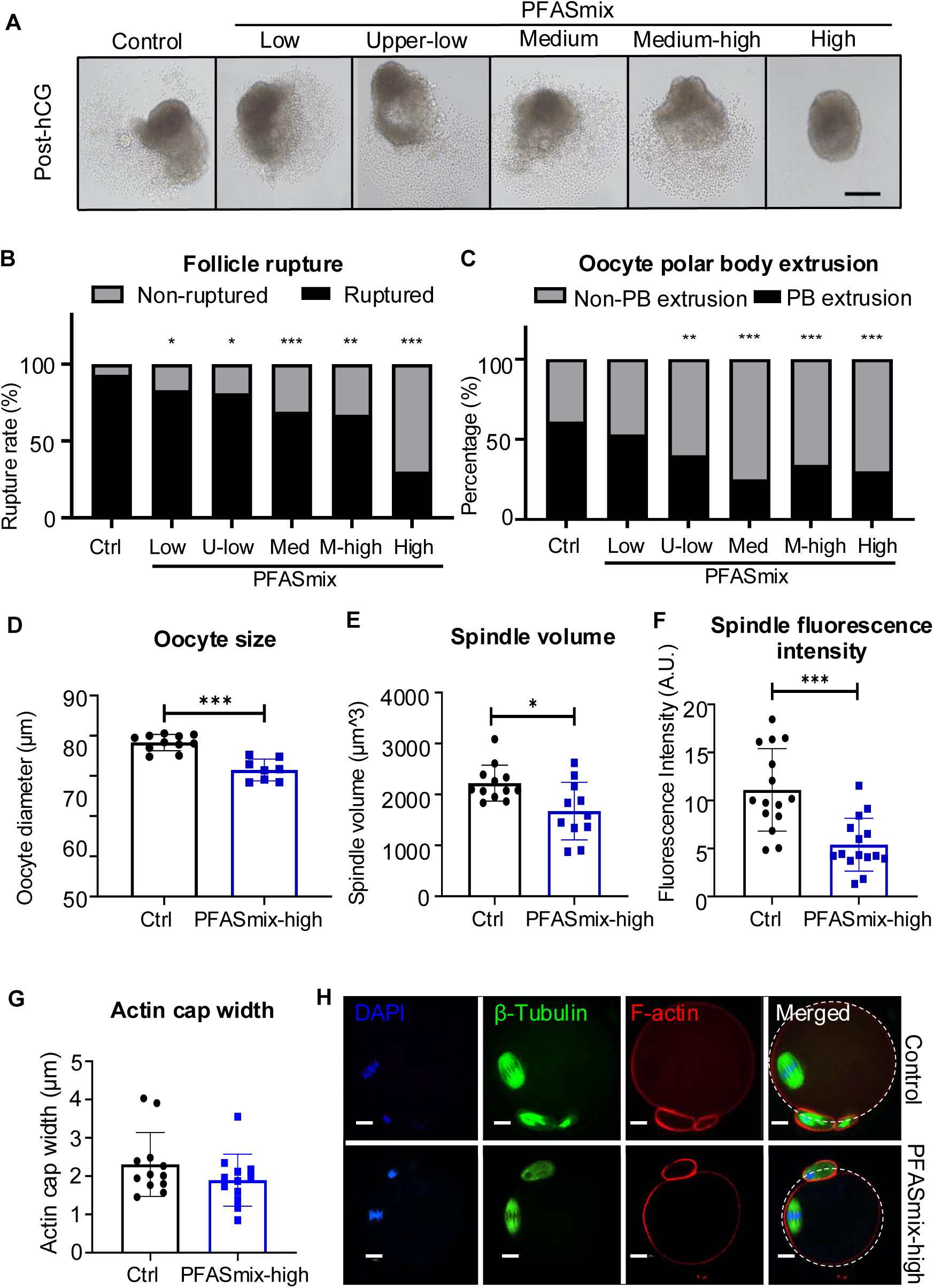
*In vitro* follicle rupture and oocyte maturation after PFAS mixture exposure in the IVFG system. (A) Representative brightfield images of ruptured follicles after human chorionic gonadotropin (hCG) induction, for control (Ctrl), low, upper-lower (U-low), medium (Med), medium-high (Med-high), and high PFAS mixture concentration groups. Scale bars: 200 µm. (B) Rate of follicle rupture and (C) percentage of collected oocytes classified as having undergone PB (polar body) extrusion or non-PB extrusion for all six treatment groups. (D) Oocyte diameter (µm) in control and PFASmix-high groups. (E) Spindle volume and (F) fluorescence intensity (A.U., arbitrary units) measured from β-tubulin immunofluorescence in control and H oocytes. (G) Actin cap width (µm). (H) Representative confocal images of *in vitro* ovulated oocytes from the control (top row) and PFASmix-high (bottom row) groups, showing DAPI, tubulin and actin stain, as well as the merged channel; dashed outlines in the merged panels indicate the oocyte membrane. Scale bars: 10 µm. *\*p<0.05, **p<0.01 and ***p<0.001*.

Oocytes recovered from ruptured follicles following PFASmix exposure were examined for maturation and cytoskeletal characteristics. Oocytes from follicles treated with PFASmix-high had significantly smaller oocytes (*p < 0.0001*, Figure 6D). Among oocytes that underwent polar body extrusion, exposure to PFASmix-high significantly reduced spindle volume (*p=0.0103*, Figure 6E, 6H) and spindle fluorescence intensity (*p=0.0002*, Figure 6F, 6H), whereas the actin-cap width was not significantly altered (*p=0.2189*; Figure 6G, 6H).

### PFASmix did not alter ovarian secretion of E2 and T *in vitro*

We examined the effects of PFASmix on steroid hormone production by measuring E2 and T in conditioned medium on day 6 of IVFG. The results showed that exposure to PFASmix did not alter E2 or T secretion on day 6 of IVFG (Figure 7A-7B).

**Figure 7.**
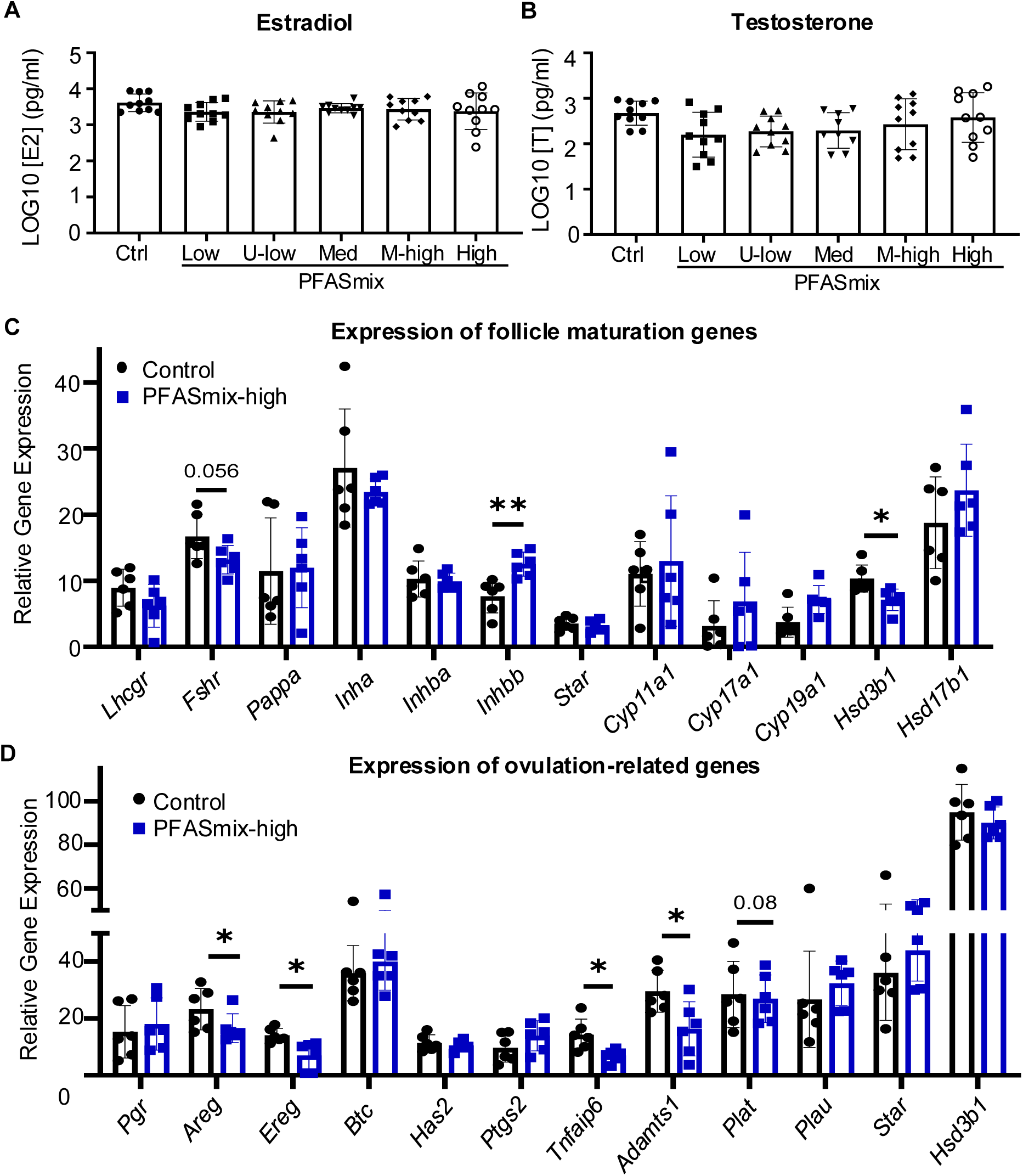
*In vitro* follicular steroid secretion and single-follicle gene expression after PFAS mixture exposure. (A) Log_₁₀_-transformed estradiol [E2] and (B) testosterone [T] concentration (pg/ml) in conditioned media collected on day 6 of IVFG for the control (Ctrl), low, upper-low (U-low), medium (Med), medium-high (M-high), and high PFAS mixture concentration groups. (C) Relative expression of follicle maturation and steroidogenic genes measured by RT-qPCR in single follicles collected on day 6 of culture from the control and PFASmix-high groups. (D) Relative expression of ovulation-related genes. *\*p<0.05, **p<0.01*.

### PFASmix-high altered the expression of genes crucial for follicle development, steroidogenesis, and ovulation *in vitro*

We next collected follicles treated with vehicle or PFASmix-high on day 6 of IVFG to examine the expression of genes crucial for follicle development and steroidogenesis, and collected follicles treated with hCG for 4 hours to examine the expression of genes crucial for ovulation. Compared to the control group, follicles treated with PFASmix-high had significantly lower expression levels of *Hsd3b1* (*p=0.0131*) and a tendency of lower expression level of *Fshr* (*p=0.0566*), and a higher expression of *Inhbb* (*p=0.0034*, Figure 7C). At 4 hours post-hCG treatment, follicles treated with PFASmix-high had significantly decreased expression of several ovulation markers, including *Areg* (*p=0.0411*), *Ereg* (*p=0.0101), Tnfaip6* (p=0.0120), and *Adamts1* (*p=0.0226*) (Figure 7D).

### PFASmix-high altered follicular cell luteinization and CL function *in vitro*

Following hCG-induced ovulation, follicular somatic cells were cultured for an additional 48 h to induce luteinization in the continued presence of PFASmix-high. PFASmix-high exposure significantly reduced P4 secretion on day 9 (p = 0.0003; Figure 8A), and reduced luteal spheroid diameter (Figure 8A-8B). Histological analysis further showed fewer cells per 1,000 µm² (p < 0.0001, Fig. 8C-8D) and an increase in TUNEL-positive cells (p = 0.0283) in PFASmix-high-treated luteal spheroids (Figure 8E, 8G). RT-qPCR further demonstrated decreased expression of the steroidogenic genes, including *Star* (*p=0.0029*), *Cyp11a1* (*p=0.0011*), and *Hsd3b1* (*p < 0.0001*), as well as *Lhcgr* (*p=0.0499*) and *Pcna* (*p=0.0144*) (Figure 8H). Reduced *Star* and *Lhcgr* expression was also visualized by RNAscope in luteal spheroids (Figure 8F).

**Figure 8.**
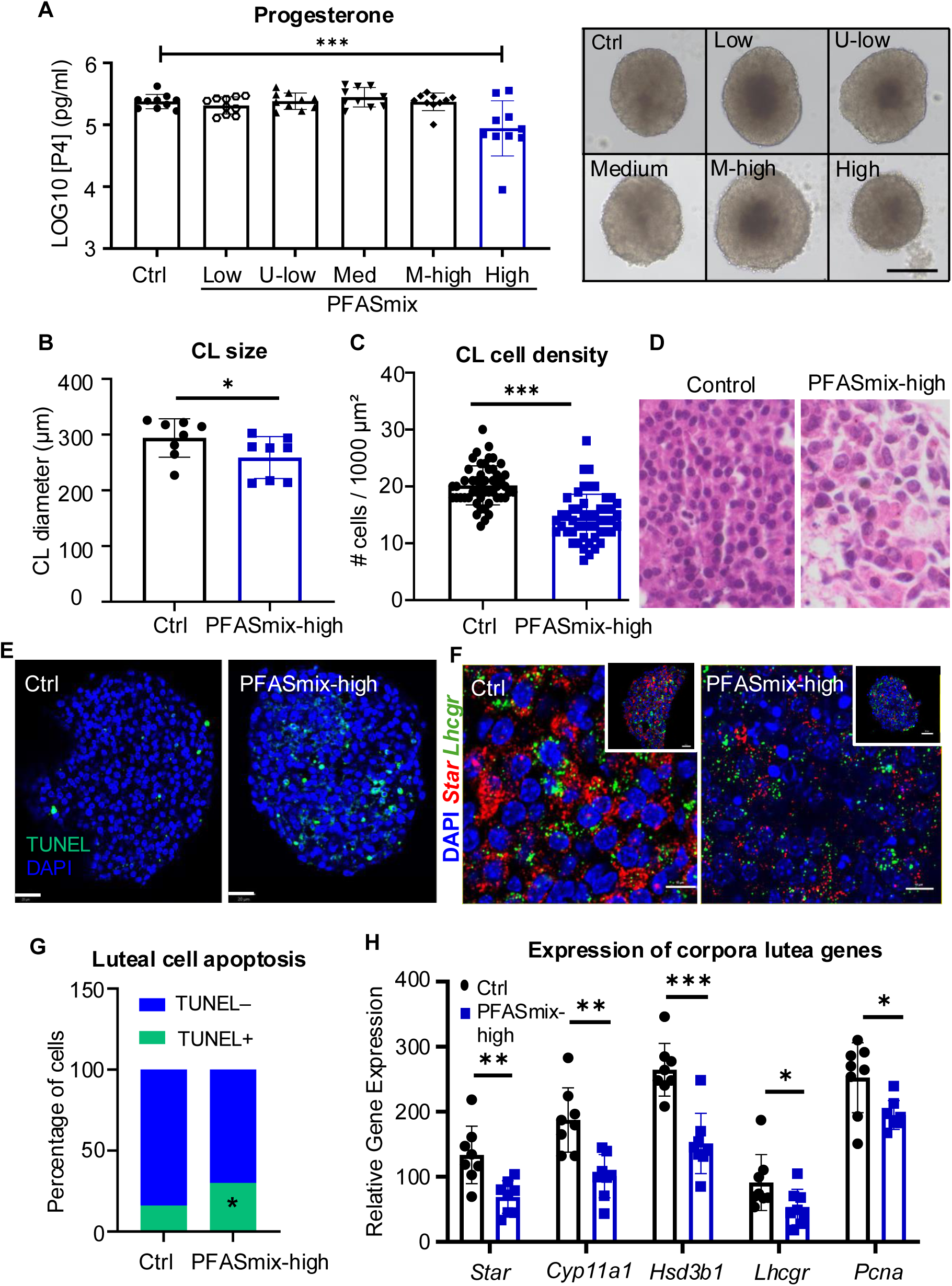
*In vitro* corpus luteum formation, progesterone secretion, morphology, and gene expression after PFAS mixture exposure. (A) Log_₁₀_-transformed progesterone [P4] concentration (pg/ml) in conditioned media collected on day 9 of culture for the control (Ctrl), low, upper-low (U-low), medium, medium-high (M-high), and high PFAS mixture concentration groups. Representative brightfield images of *in vitro*–formed CL (corpora lutea). Scale bar: 100 µm. (B) CL diameter (µm) in the control and PFASmix-high groups. (C) Luteal cell density, expressed as the number of cells per 1000 µm². (D) Representative photomicrograph of CL stained with hematoxylin/eosin, showing the histoarchitecture in control and PFASmix-high groups. (E) Representative TUNEL images of *in vitro*–formed CL from the control and PFASmix-high groups, counterstained with DAPI. (F) Representative RNAscope in situ hybridization images from the control and PFASmix-high group, showing DAPI (blue), *Star* (red), and *Lhcgr* (green). Scale bar:10 µm. (G) Percentage of luteal cells scored as TUNEL-negative or TUNEL-positive. Scale bar: 20 µm. (H) Relative expression of steroidogenic enzymes, LH receptor, and a cell proliferation marker (*Pcna*), on CL from control and PFASmix-high groups. *p<0.05, **p<0.01, *and ***p<0.001*.

### PFASmix altered PPAR**γ**-associated gene expression in ovarian follicles *in vitro*

Based on our previous findings implicating PPARγ signaling in the ovarian effects of individual long-chain PFAS, we examined whether direct PFASmix exposure altered the expression of PPARγ-associated genes in the 3D follicle culture model. The results showed that exposure to PFASmix-high significantly increased the expression of *Adipoq* (*p=0.0296*), *Serpine1* (*p=0.0299*), *Txnip* (*p=0.0161*), and *Ucp1* (*p=0.0438*) in grown antral follicles on day 6 of IVFG (Figure 9), indicating that direct PFASmix exposure is associated with altered PPARγ-related transcriptional responses in ovarian follicles.

**Figure 9.**
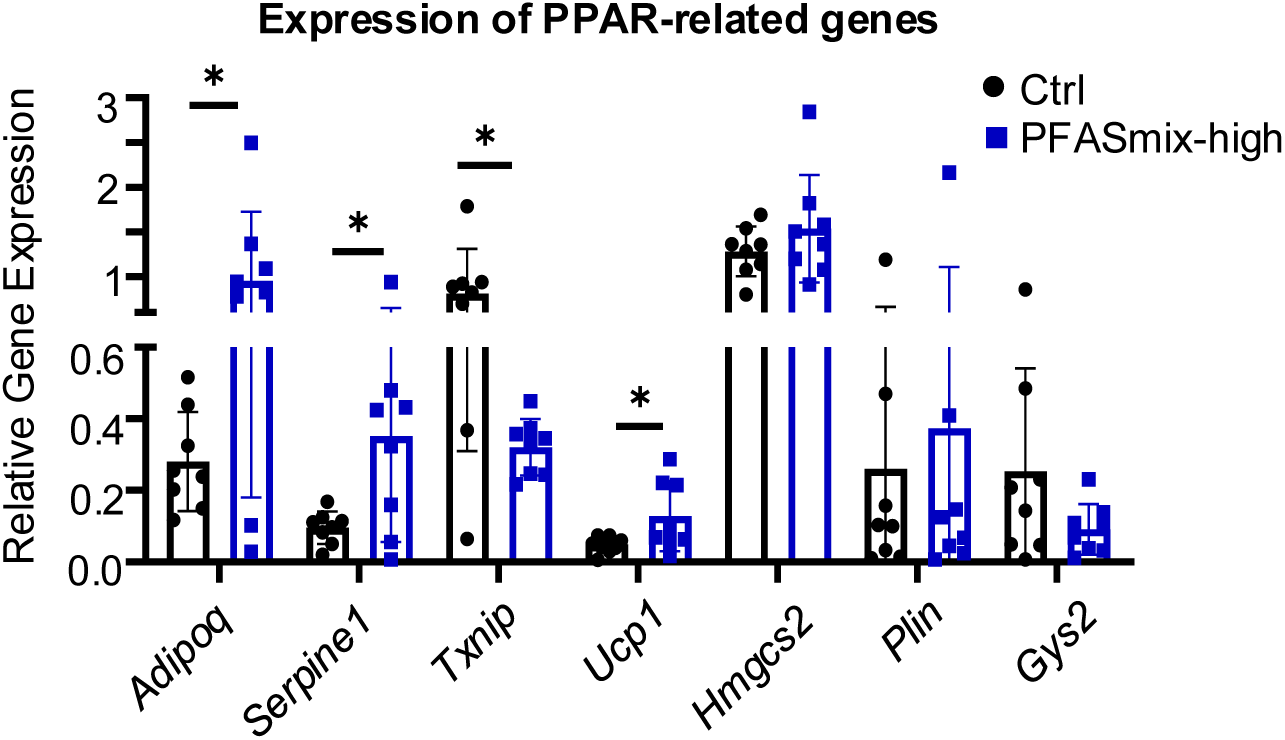
Relative expression of PPAR-(peroxisome proliferator-activated receptor-) related genes in follicles after *in vitro* PFAS mixture exposure. *\*p<0.05*.

### Benchmark dose/concentration modeling identifies sensitive reproductive endpoints

BMD modeling was used to quantitatively compare the dose or concentration-response relationships across selected reproductive endpoints. Among the *in vivo* outcomes, the number of complete estrous cycles was the most sensitive endpoint, with a BMD of 4.89 × 10⁻□mg/mL (BMDL, 3.46 × 10⁻□mg/mL; BMCU, 8.67 × 10⁻□mg/mL). In the 3D follicle culture model, follicle rupture was the most sensitive endpoint, with a BMC of 5.00 × 10⁻³ mg/mL, corresponding to approximately 10.8 µM (BMCL, 2.00 × 10⁻³ mg/mL [4.3 µM]; BMCU, 8.80 × 10⁻² mg/mL [189 µM]).

## Discussion

In the current study, we used complementary *in vivo* mouse exposure model and *in vitro* 3D ovarian follicle culture system to determine how the mixture of 5 commonly detected long-chain PFAS affects female ovarian functions. The results revealed that PFASmix exposure produced a coordinated pattern of reproductive toxicity spanning estrous cyclicity, follicle development, ovulation, oocyte maturation, and luteal development and function. *In vivo*, PFASmix at PFAS contamination sites or occupational exposure levels disrupted mouse estrous cyclicity, altered circulating E2 and T, increased follicular atresia, and impaired oocyte maturation. Direct exposure of intact follicles *in vitro* recapitulated several of these effects and further revealed concentration-dependent impairment of follicle growth and ovulation, disruption of periovulatory gene expression, and compromised luteinization and P4 production. Together, these findings extend our previous studies of individual PFAS (42) by demonstrating that exposure to a long-chain PFAS mixture can directly disrupt multiple sequential stages of ovarian functions.

The disruption of estrous cyclicity and reproductive hormones observed *in vivo* is consistent with epidemiological evidence linking PFAS exposure to altered menstrual cyclicity and reproductive endocrine function (35, 36, 56–64). Exposure to PFASmix-high prolonged diestrus, reduced time in estrus and the number of complete cycles, decreased circulating E2, and increased T. Although PFAS has been shown to disrupt hypothalamic-pituitary signaling in experimental models (41), FSH was unchanged, and LH showed only a nonsignificant decrease in our study. However, because gonadotropins were measured at a single time point, these results cannot exclude effects on pulsatile secretion or the preovulatory LH surge. Importantly, the *in vitro* follicle culture experiments showed impaired follicular function in the absence of the hypothalamic-pituitary axis, supporting the notion that the ovary is a direct target of PFASmix toxicity.

The gonadotropin-dependent phase of follicle development appeared particularly sensitive to PFASmix exposure. *In vivo*, exposure to PFASmix-high increased antral follicular atresia and granulosa cell apoptosis without significantly altering the numbers of follicles at individual developmental stages. Large antral follicles from PFASmix-exposed mice also showed decreased *Lhcgr* and increased *Amh* expression, a pattern consistent with impaired acquisition of differentiated preovulatory follicular phenotype. Similarly, direct PFASmix exposure inhibited follicle growth *in vitro* and reduced *Hsd3b1* expression. These findings are consistent with our previous study that individual long-chain PFAS impair follicle development and steroidogenesis (42), suggesting that granulosa cell survival and differentiation are important targets of long-chain PFASmix.

Exposure to PFASmix also disrupted the transition from follicle development to ovulation. *In vitro* follicle rupture and oocyte polar body extrusion decreased in a concentration-dependent manner, accompanied by suppression of *Areg, Ereg, Tnfaip6,* and *Adamts1* following hCG stimulation. These genes participate in the coordinated periovulatory program required for cumulus expansion, extracellular-matrix (ECM) remodeling, and follicular rupture (65). Their suppression, together with reduced *Lhcgr* in antral follicles from PFASmix-exposed mice, suggests that exposure to PFASmix interferes with the ability of follicles to respond appropriately to the ovulatory stimulus. Oocyte maturation was also affected in both *in vivo* and *in vitro* models. Oocytes from exposed mice and PFASmix-treated follicles were smaller, showed reduced polar body extrusion, and exhibited alterations in meiotic spindle characteristics. These findings are consistent with previous evidence that PFAS can disrupt meiotic spindle organization and actin dynamics in oocytes (4, 31, 32). Because fertilization and embryo development were not evaluated, these effects are best interpreted as impaired oocyte maturation and cytoskeletal organization rather than reduced developmental competence.

A notable finding was that PFASmix toxicity extended beyond ovulation to CL development and function. Following hCG-induced ovulation *in vitro*, PFASmix-exposed follicles formed smaller CL spheroids with fewer luteal cells and increased apoptosis. These structural changes were accompanied by decreased the expression of *Star, Cyp11a1,* and *Hsd3b1* and reduced hormonal P4 secretion, indicating impaired luteal steroidogenesis. *Lhcgr* also decreased, suggesting compromised responsiveness to luteotropic LH/hCG signaling. Thus, exposure to PFASmix disrupted not only gonadotropin-dependent follicular development and ovulation but also the subsequent follicle-to-CL transition. Given the essential role of luteal P4 in establishing and maintaining early pregnancy, particularly uterine receptivity and embryo implantation, luteal dysfunction may represent an important but understudied reproductive consequence of PFAS exposure.

Our findings also support a potential involvement of PPARγ-associated signaling. We previously identified PPARγ activation as a molecular initiating event (MIE) for PFNA-induced ovarian toxicity and showed that pharmacological inhibition of PPARγ reversed PFNA-induced molecular and functional defects (42). Here, the direct PFASmix exposure increased the expression of several genes associated with the PPARγ signaling, including *Adipoq, Serpine1, Txnip,* and *Ucp1*, in cultured follicles. These findings suggest that PPARγ-associated signaling may represent a convergent ovarian response to exposure to a long-chain PFASmix. However, because PPARγ activity was not directly manipulated in the present study, it remains to be determined whether PPARγ is the bona fide mediator of the ovarian response to direct exposure to long-chain PFAS mix.

Another strength of this study is the integration of environmentally and biologically informed mixture designs with quantitative dose and concentration-response analysis. The *in vivo* PFASmix was derived from documented contaminated groundwater, with a concentration range spanning public drinking-water and highly contaminated groundwater scenarios (46, 47, 66, 67). The *in vitro* mixture composition was based on human serum PFAS profiles, with concentrations extending from general population levels toward those reported in occupationally exposed populations (5, 49–52, 68–70). Direct equivalence among drinking water, serum, and *in vitro* follicle culture medium concentrations should not be assumed, as PFAS toxicokinetics, protein binding, tissue distribution, and exposure duration differ across these settings. Nevertheless, these complementary designs enabled evaluation across environmentally informed external exposures and biomonitoring-informed internal concentrations. BMD modeling further identified the number of complete estrous cycles and follicle rupture as sensitive *in vivo* and *in vitro* endpoints, respectively, providing quantitative points of departure (POD) for PFASmix-induced reproductive toxicity.

There are several limitations we need to consider in this study. Internal PFAS concentrations in the blood and follicular fluid were not measured, limiting direct comparison with human biomonitoring data. Molecular analyses were primarily conducted at the highest concentration, and the causal contribution of PPARγ to mixture toxicity was not directly tested. In addition, the PFASmix contained five legacy long-chain PFAS and therefore does not capture emerging long- and short-chain PFAS. Finally, species differences in PFAS toxicokinetics and PPAR signaling should be considered when extrapolating these findings to humans (4, 71).

In conclusion, our results demonstrate that exposure to an environmentally and human-relevant long-chain PFASmix disrupted multiple sequential stages of ovarian function, including gonadotropin-dependent follicle development and survival, ovulation, oocyte maturation, and luteal development and steroidogenesis. The concordance between the *in vivo* and *in vitro* follicle culture models supports a direct ovarian impact of PFASmix, while BMD modeling identifies estrous cyclicity and ovulation as sensitive female reproductive endpoints. These findings extend the evidence from individual PFAS to a complex mixture of PFAS and provide mechanistic and quantitative evidence relevant to female reproductive health.

## Authors’ roles

A.R.R. Caceres contributed to experimental design, data collection and analysis, and manuscript writing. J. Zhang contributed to the data analysis and manuscript writing. T. Costanza contributed to histological staining and TUNEL staining. G. Guo contributed to the data interpretation and manuscript editing. S. Xiao and Q. Zhang received the funding support, conceived the project, designed experiments, analyzed and interpreted data, wrote the manuscript, and provided final approval of the manuscript.

## Funding and acknowledgments

This work was supported by the National Institutes of Health (NIH) R01ES032144 and R01ES035766. This work was also partially supported by the Assistant Secretary of Defense for Health Affairs through the Toxic Exposures Research Program (TERP), endorsed by the Department of Defense (DOD) under Award No. HT9425-23-1-0809. Opinions, interpretations, conclusions, and recommendations are those of the author and are not necessarily endorsed by the DOD.

## Notes

### Competing Interest Statement

The authors have declared no competing interest.

