## Supplemental Table S1 to S7 for "Exposure to a mixture of long-chain per- and polyfluoroalkyl substances (PFAS) disrupts ovarian follicle development, ovulation, and luteinization in female mice"

**Table of contents**

**Table S1.** List of PFAS used in the current study.

**Table S2.** PFAS concentrations in Marine Corps Base Camp Lejeune, North Carolina

**Table S3.** *In vivo* PFAS mixture concentration summary.

**Table S4.** Serum PFAS data extracted from NHANES(1) corresponding to the US population.

**Table S5.** *In vitro* PFAS mixture concentration summary.

**Table S6.** Specifications of the ELISA kits used for measuring serum hormones.

**Table S7.** Primer sequences of selected genes used for qPCR experiments.

**Table S1.** List of PFAS used in the current study.

| ***PFAS*** | ***Classification*** | ***Functional group*** | ***CAS#*** | ***Purity*** |
| --- | --- | --- | --- | --- |
| **PFOS** | LEGACY | PFSAs | 77282 | ≥95% |
| **PFHxS** | LEGACY | PFSAs | 50929 | ≥98% |
| **PFDA/PFDeA** | EMERGING | PFCAs | 177741 | ≥98% |
| ***PFNA*** | LEGACY | PFCAs | 394459 | ≥97% |
| **PFOA** | LEGACY | PFCAs | 171468 | ≥95% |

PFSAs: perfluoroalkyl sulfonic acids (CnF2n+1 - SO3H), PFCAs: perfluoroalkyl carboxylic acids (CnF2n+1 – COOH). PFOA (Perfluorooctanoic acid), PFOS (Hepta-deca-fluorooctanesulfonic acid potassium salt), PFNA (Perfluorononanoic acid), PFHXs (Tridecafluorohexane-1-sulfonic acid potassium salt), and PFDA (Perfluorodecanoic acid) were purchased from Sigma-Aldrich (St. Louis, MO).

**Table S2.** PFAS concentrations in Marine Corps Base Camp Lejeune, North Carolina

| *PFAS compound* | Camp Lejeune, NC (USMC Base; AFFF fire training groundwater)(2) |
| --- | --- |
| *PFHxS* | 101,000 |
| *PFNA* | 206 |
| *PFOS* | 40,461 |
| *PFOA* | 25,100 |
| *PFDA* | 72 |
| TOTAL PFAS (sum) | ~167,000 ng/L |

All concentrations are in ng/L (= ppt). USMC: United States Marine Corps. AFFF: Aqueous film-forming foam.

**Table S3.** *In vivo* PFAS mixture concentration summary.

| **PFASmix** | **Low mg/ml** | **Medium mg/ml** | **Medium-high mg/ml** | **High mg/ml** |
| --- | --- | --- | --- | --- |
| **PFHxS** | 4.70E-08 | 4.71E-06 | 4.71E-05 | 4.71E-04 |
| **PFNA** | 1.65E-09 | 1.65E-07 | 1.65E-06 | 1.65E-05 |
| **PFOS** | 3.50E-07 | 3.51E-05 | 3.51E-04 | 3.51E-03 |
| **PFOA** | 3.45E-08 | 3.46E-06 | 3.46E-05 | 3.46E-04 |
| **PFDA** | 7.14E-10 | 7.16E-08 | 7.16E-07 | 7.16E-06 |
| **TOTAL mg/ml** | **4.34E-07** | **4.35E-05** | **4.35E-04** | **4.35E-03** |

**Table S4.** Serum PFAS data extracted from NHANES (1) corresponding to the US population.

| PFAS abbreviation | Chemical name | LLOD ng/ml | Mean ng/ml | Median ng/ml | 95th percentile |
| --- | --- | --- | --- | --- | --- |
| PFOS* | Perfluorooctane sulfonic acid | 0.1 | 6.111 | 3.8 | 17.8 |
| PFOA | Perfluorooctanoic acid | 0.1 | 1.689 | 1.37 | 3.77 |
| PFDeA | Perfluorodecanoic acid | 0.1 | 0.2756 | 0.2 | 0.8 |
| PFNA | Perfluorononanoic acid | 0.1 | 0.6469 | 0.5 | 1.6 |
| PFHxS | Perfluorohexane sulfonic acid | 0.1 | 1.567 | 1 | 4 |

Data source: National Health and Nutrition Examination Survey 2017-2020.

*Total PFOS value was obtained after the sum of linear + branched isomers as reported by NHANES.

**Table S5.** *In vitro* PFAS mixture concentration summary.

| **PFAS MIX** | **Low mg/ml** | **Upper-low mg/ml** | **Medium mg/ml** | **Medium-high mg/ml** | **High mg/ml** |
| --- | --- | --- | --- | --- | --- |
| **PFHXS** | 1.43E-06 | 1.43E-05 | 1.43E-04 | 1.43E-03 | 1.43E-02 |
| **PFNA** | 6.45E-07 | 6.45E-06 | 6.45E-05 | 6.45E-04 | 6.45E-03 |
| **PFOS** | 5.71E-06 | 5.71E-05 | 5.71E-04 | 5.71E-03 | 5.71E-02 |
| **PFOA** | 1.69E-06 | 1.69E-05 | 1.69E-04 | 1.69E-03 | 1.69E-02 |
| **PFDA** | 2.73E-07 | 2.73E-06 | 2.73E-05 | 2.73E-04 | 2.73E-03 |
| **TOTAL** | **9.75E-06** | **9.75E-05** | **9.75E-04** | **9.75E-03** | **9.75E-02** |

**Table S6.** ELISA kits used for measuring serum hormones.

| **Hormone** | **Sensitivity** | **Range** | **Intra-assay %CV** | **Inter-assay %CV** |
| --- | --- | --- | --- | --- |
| **FSH** | 0.016 ng/ml | 0.016 - 8.0 ng/ml | 3.6 | 5.1 |
| **LH** | 0.016 ng/ml | 0.016 - 4.0 ng/ml | 2.2 | 7.3 |
| **AMH** | 0.2 ng/mL | 0.2 – 15 ng/mL | 2.1 | 7.6 |
| **Inhibin-B** | 8 pg/mL | 8 - 942 pg/mL | 5.1 | 7.9 |
| **E2** | 5 pg/mL | 5 – 3,200 pg/mL | 4.3 | 6.2 |
| **T** | 10 ng/dL | 10 – 1,600 ng/dL | 3.6 | 4.7 |
| **P4** | 0.15 ng/ml | 0.15 - 40.0 ng/ml | 3.1 | 8.1 |

**Table S7.** Primer sequences of selected genes used for qPCR experiments.

| Gene | Forward primer | Reverse primer |
| --- | --- | --- |
| *Actb* | 5'-GTGACGTTGACATCCGTAAAGA-3' | 5'-GCCGGACTCATCGTACTCC-3' |
| *Fshr* | 5'-GAGGCAGATGTCTTCTCCAAC-3' | 5'-TCGGACACTGGGAAGATTCTGG-3' |
| *Lhcgr* | 5'-GACGCTAATCTCGCTGGAGT-3' | 5'-GGCCTGCAATTTGGTGGAAG-3' |
| *Amh* | 5'-CAGGCCCTGTTAGTGCTATACCCT-3' | 5'-GAAGTCCACGGTTAGCACCAAA-3' |
| *Pappa* | 5'-CAGAAAGCCAGCACCTGTAGCT-3' | 5'-GGCAAAGGTCACATGCTGATCC-3' |
| *Inha* | 5'-CAGGCTATCCTTTTCCCAGCTAC-3' | 5'-AAGTCACCTGGTGGCTGCGTAT-3' |
| *Inhba* | 5'-GGAGATAGAGGACGACATTGGC-3' | 5'-ACGCTCCACTACTGACAGGTCA-3' |
| *Inhbb* | 5'-CTCCGAGATCATCAGCTTTGCAG-3' | 5'-GGAGCAGTTTCAGGTACAGCCA-3' |
| *Comp* | 5'-GTGCCCAACTTTGACCAGAGTG-3' | 5'-ACAGGCATCACCCACAAAGTCG-3' |
| *Star* | 5'-TTGGGCATACTCAACAACCA-3' | 5'-CCTTGACATTTGGGTTCCAC-3' |
| *Cyp17a1* | 5'-CCAGGACCCAAGTGTGTTCT-3' | 5'-CCTGATACGAAGCACTTCTCG-3' |
| *Cyp19a1* | 5'-CATGGTCCCGGAAACTGTGA-3' | 5'-GTAGTAGTTGCAGGCACTTC-3' |
| *Cyp11a1* | 5'-TCAAAGCCAGCATCAAGGAGA-3' | 5'-TGGCAAAGCTAGCCACCTGTA-3' |
| *Hsd3b1* | 5'-AGTGATGGAAAAAGGGCAGGT-3' | 5'-GCAAGTTTGTGAGTGGGTTAG-3' |
| *Hsd17b1* | 5'-ACTGTGCCAGCAAGTTTGCG-3' | 5'-AAGCGGTTCGTGGAGAAGTAG-3' |
| *Esr1* | 5'-GCGCAAGTGTTACGAAGTG-3' | 5'-TTCGGCCTTCCAAGTCATC-3' |
| *Esr2* | 5'-GGTCCTGTGAAGGATGTAAGGC-3' | 5'-TAACACTTGCGAAGTCGGCAGG-3' |
| *Pgr* | 5'-ATGGTCCTTGGAGGTCGTAA-3' | 5'-GTGGCGGGACCAGTTGAAT-3' |
| *Areg* | 5'-GCTGAGGACAATGCAGGGTAA-3' | 5'-AGTGACAACTGGGCATCTGG-3' |
| *Ereg* | 5'-GCATCCCAGGAGAATCCGAG-3' | 5'-GTGTAGCCCACTTCACATCTGC-3' |
| *Btc* | 5'-AAACCCACTTCTCTCGGTGC-3' | 5'-GCCTTTCTCACAGATGCAGG-3' |
| *Has2* | 5'-GCCGGTCGTCTCAAATTCATC-3' | 5'-ACCTCTCACAATGCATCTTGTTC-3' |
| *Ptgs2* | 5'-TGGGGGAAGAAATGTGCCAA-3' | 5'-CAGCCATTTCCTTCTCTCCTGT-3' |
| *Tnfaip6* | 5'-GCTACAACCCACATGCAAAGG-3' | 5'-CTGACCGTACTTGAGCCGAA-3' |
| *Adamts1* | 5'-TTGAATGGTGTGAGTGGCGA-3' | 5'-CCATCAAACATTCCCCGTGTC-3' |
| *Plat* | 5'-AAGAAGCAAGCACTCTCGGG-3' | 5'-TCATCTCTGCAGGTCGCTCT-3' |
| *Plau* | 5'-CATCCAGTCCTTGCGTGTCT-3' | 5'-CCAAGTACACTGCCACCTTCA-3' |
| *Pcna* | 5'-CAAGTGGAGAGCTTGGCAATGG-3' | 5'-GCAAACGTTAGGTGAACAGGCTC-3' |
| *Fabp1* | 5'-AGTCAAGGCAGTCG-3' | 5'-ATGTCGCCCAATGTC-3' |
| *Cyp4a10* | 5'-CTCCAAGTGGCCTCT-3' | 5'-TAAGTAGGCCTTGC-3' |
| *Acox1* | 5'-CAAAGAAACCCCTC-3' | 5'-GCTGTGTGCCGTCT-3' |
| *Gsta* | 5'-CGCCACCAAATATG-3' | 5'-TTGCCCAATCATTTC-3' |
| *Cyp2b10* | 5'-GACTTTGGGATGGG-3' | 5'-CCAAACACAATGGA-3' |
| *Akr1b7* | 5'-AAGCGGGAGGATCT-3' | 5'-TCAGATCCGAGAGG-3' |
| *Cyp3a11* | 5'-ACAAGGGTTTATGG-3' | 5'-GTCTGTGACAGCAA-3' |
| *Cyp2b9* | 5'-TGGCCACCATGAAA-3' | 5'-GCTGTGATGCACTG-3' |
| *Cyp17a1* | 5'-CCAGGACCCAAGTGTGTTCT-3' | 5'-CCTGATACGAAGCACTTCTCG-3' |
| *Adipoq* | 5'-TGTTCCTCTTAATCCTGCCCA-3' | 5'-CCAACCTGCACAAGTTCCCTT-3' |
| *Serpine1* | 5'-TCTGGGAAAGGGTTCACTTTACC-3' | 5'-GACACGCCATAGGGAGAGAAG-3' |
| *Txnip* | 5'-GGCCGGACGGGTAATAGTG-3' | 5'-AGCGCAAGTAGTCCAAAGTCT-3' |
| *Ucp1* | 5'-GGACAGTACCCAAGCGTACCA-3' | 5'-CGCAGAAAAGAAGCCACAAAC-3' |
| *Hmgcs2* | 5'-AGAGAGCGATGCAGGAAACTT-3' | 5'-AAGGATGCCCACATCTTTTGG-3' |
| *Plin* | 5'-CTGTGTGCAATGCCTATGAGA-3' | 5'-CTGGAGGGTATTGAAGAGCCG-3' |
| *Gys2* | 5'-CGCTCCTTGTCGGTGACATC-3' | 5'-CATCGGCTGTCGTTTTGGC-3' |
